# Essential role for Ggct in erythrocyte antioxidant defense

**DOI:** 10.1101/2021.06.21.449332

**Authors:** Zaoke He, Xiaoqin Sun, Shixiang Wang, Dongsheng Bai, Xiangyu Zhao, Ying Han, Piliang Hao, Xue-Song Liu

**Affiliations:** School of Life Science and Technology, ShanghaiTech University, Shanghai 201203, China; Shanghai Institute of Biochemistry and Cell Biology, Chinese Academy of Sciences, Shanghai, China; University of Chinese Academy of Sciences, Beijing, China

**Keywords:** GGCT, ROS, anemia, glutathione, red blood cell, mouse model

## Abstract

*GGCT* encodes γ-glutamyl cyclotransferase enzyme activity, and its expression is up-regulated in various human cancers. γ-glutamyl cyclotransferase enzyme activity was originally purified from human red blood cells (RBCs), however physiological function of *GGCT* in RBCs is still not clear. Here we reported that *Ggct* deletion in mouse leads to splenomegaly and progressive anemia phenotypes, due to elevated oxidative damage and shortened life span of *Ggct*^−/−^ RBCs. *Ggct^−/−^* RBCs have increased reactive oxygen species (ROS), and are more sensitive to H_2_O_2_ induced damage compared to control RBCs. Glutathione (GSH) and GSH synthesis precursor L-cysteine are decreased in *Ggct^−/−^* RBCs. Our study suggests a critical function of Ggct in RBC redox balance and life span maintenance through regulating GSH metabolism.

## Background

GGCT (γ-glutamyl cyclotransferase) was also named as C7orf24, and was reported to be up-regulated in various human cancers, including bladder urothelial carcinoma (Kageyama*, et al* 2007), breast cancer (Gromov*, et al* 2010), lung, esophagus, stomach, bile duct and uterine cervix cancer (Amano*, et al* 2012). In 2008, Oakley et al. cloned the cDNA encoding human γ-glutamyl cyclotransferase enzyme activity from human red blood cells (RBCs), and this study identified C7orf24 as GGCT, an enzyme in the γ-glutamyl cycle (Oakley, et al 2008). γ-glutamyl cyclotransferase catalyzes the following reaction: γ-glutamyl–amino acid ⟶ 5-oxoproline + amino acid. The physiological function of this enzyme activity is not clear. Meister proposed that this enzyme is a critical component of γ-glutamyl cycle, and is involved in glutathione (GSH) degradation and amino acid transport through plasma membrane (Meister 1974). In γ-glutamyl cycle, extracellular GSH can be hydrolyzed by membrane-bound γ-glutamyl transpeptidase (GGT) to cysteinyl-glycine and γ-glutamyl– amino acid dipeptide (Anderson 1998, Meister 1988). In the cytoplasm, γ-glutamyl cyclotransferase cleaves the γ-glutamyl–amino acid to give 5-oxoproline and amino acid (Meister 1974). Currently, the specific physiological function of γ-glutamyl cycle has been debated (Bachhawat and Yadav 2018). Recent studies also identified GSH cytoplasmic degradation pathway through ChaC family proteins, ChaC family of proteins function as γ-glutamyl cyclotransferases acting specifically to degrade glutathione but not other γ-glutamyl peptides (Kumar*, et al* 2012). Till now, the physiological function of GGCT in mammals is still not clear.

To study the physiological function of GGCT, we reported the generation of the first *Ggct* knockout mouse, *Ggct* deletion is compatible with normal mouse embryonic development (He*, et al* 2019). Here we show that in young ages, *Ggct^−/−^* mice appear normal, while *Ggct^−/−^* adult or old mice show splenomegaly, progressive anemia and lack of activities phenotypes. GGCT enzyme was originally purified from human RBCs, however the function of GGCT in RBC is still unknown. In the present study, we examined the *in vivo* function of Ggct in RBC of *Ggct^−/−^* mice, and found that Ggct plays a critical role in protecting RBC against oxidative stress in mice.

## Materials and Methods

### Western blot

Spleen tissue was weighed and wash three times with cold PBS. Cell lysate buffer (20Mm HEPES PH=7.5; 150Mm NaCl; 1%NP40) was added to spleen tissue and crushed by Homogenizer. The homogenate was passed through a 40 μm filter, and boiled with protein loading buffer. Anti-GGCT antibody (ab198503, Abcam), Anti-β-actin antibody (AC-15, Sigma), were used for Western blot. For chemiluminescence, horseradish peroxidase-conjugated secondary antibodies and Western Lightining^®^ Plus-ECL (NEL105001EA, PerkinElmer) were used.

### *Ggct^−/−^* mouse genotyping with PCR

Genomic DNA was obtained by the Mouse Direct PCR Kit (Bimake, B40013, Shanghai) from mouse tail. *Ggct^−/−^* mice were genotyped with the following primers:

GGCT-KO-F: 5’-TGAGTCATAGATCTGACAGCAAGAG-3’
GGCT-KO-R: 5’-ATAACCCCTGTGAACCATCATTCA-3’

Predicted PCR product size for wild type allele is 994bp, *Ggct^−/−^* allele is 382 bp. *Ggct^−/−^* mouse lines were generated by Shanghai Model Organisms Center, Inc. (SMOC). All mouse studies were carried out in strict accordance with the guidelines of the Institutional Animal Care and Use Committee (IACUC) at the School of Life Science and Technology, ShanghaiTech University.

### Histological and hematological analyses

15-week-old male and female mice were weighed and sacrificed. Selected organs, including spleen, liver, kidney, heart, and lung, were removed and weighed to calculate an organ index (organ index=organ weight/body weight). For the histologic study, spleens were fixed in 4% buffered neutral formalin, embedded in paraffin, and stained with hematoxylin and eosin. Tissue section were examined with an Olympus VS120 microscope. Peripheral blood sample were harvested in EDTA-coated microtubes (IDEXX, 98-0010316-00, UK) by retro-orbital sinus bleeding and analyzed with a Procyte Dx Veterinary Hematology Analyzer (IDEXX, B6972, UK). Blood smears were stained with Giemsa and analyzed with Olympus BX53.

### Erythrocyte differentiation stage quantification by flow cytometry

Mouse spleen and bone marrow cells were mechanically dissociated through a 70um strainer and washed with cold phosphate-buffered saline containing 2% fetal calf serum. Splenocyte single cell suspensions were double-stained with antibodies against fluorescein isothiocyanate-conjugated CD71 (CD71-FITC) and phycoerythrin-conjugated erythroid antigen (Ter119-PE). Flow cytometry was performed using a FACSCalibur.

### RBCs life span determination

RBCs of wild-type or *Ggct^−/−^* mice were labelled with 4 mM CMFDA (Molecular Probes) which emits green fluorescence after cleavage by intracellular esterases. The labelled cells were injected into wild-type or *Ggct^−/−^* recipient mice intravenously. Blood was collected at indicated time points, and CMFDA labelled cells were quantified by flow cytometry.

### Osmotic fragility assay

RBCs were harvested using heparin-coated tubes, and then suspended in varying concentrations of NaCl. The samples were incubated at room temperature for 10 min and centrifuged at 1500g for 10 min to remove unlysed cells and stromal cells. The absorbance of the supernatant was measured at 540nm in a spectrophotometer (Peinado*, et al* 1992). The lyses percentage of RBCs was calculated from the absorbance, and a fragility curve was generated by plotting varying salt concentrations versus hemolysis.

### Erythropoietin (EPO) quantification by enzyme-linked immunosorbent assay

EPO level in blood plasma was determined using the mouse EPO enzyme-linked immunosorbent assay kit (Jiningshiye, A00895-2, shanghai). Heparinized blood was collected from wild-type and *Ggct^−/−^*, and then blood samples were centrifuged at 1,000g for 10 min to obtain the plasma. 50ul of plasma was taken for the experiment according to the manufacturer’s protocol.

### Metabolic cage experiment

Wild-type and *Ggct^−/−^* mice were individually housed in Oxymax Comprehensive Laboratory Animal Monitoring System (CLAMS) (Columbus Instruments, Columbus, OH, USA) sealed chambers, each of which was equipped with an O_2_ electrochemical sensor, a CO_2_ infrared sensor and infrared beam activity sensors. The airflow rate was 0.5 L/min per cage. Mice were placed in the chamber 1 day prior to the start of measurements to allow for acclimation to the new environment. The metabolic data collected include the volume of O_2_ consumed (V_O2_), volume of CO_2_ generated (V_CO2_), respiratory exchange ratio (RER) (RER = V_CO2_/V_O2_) and heat produced. O_2_ consumption and CO_2_ production were measured over a 2-min period, which was repeated every 10 min. V_O2_ and V_CO2_ values were normalized to the body weights of the mice (ml/kg/h). The infrared beam interruptions in both horizontal (X) and vertical (Z) directions were used to quantify the activity of mice. Any horizontal beam breakage was recorded as total activity count. Any vertical beam breakage was recorded as total activity count. During the recording, the mice were deprived of food, and with free access to water.

### Metabolomics mass spectrometry

Peripheral blood was harvested into heparin-coated tubes, and centrifuged at 4℃ 1000g for 10 minutes. 100 ul plasma and blood cells were taken and add 1ml of MeOH: ACN (acetonitrile) H2O (2:2:1, V/V) solvent mixture to the sample and vortex 30s, sonicate for 10 min and incubate for 1 hour at −20℃, centrifuge 15min at 13000rpm and 4℃. Take supernatant and evaporate to dryness at 4℃ using a vacuum concentrator; 100ul of ACN:H2O(1:1/V/V) sonicate 10 min, centrifuge 15min at 13000rpm and 4℃. keep supernatant in −80℃prior to LC/MS analysis. Analyses were performed using a Waters Acquity I Class UPLC system connected to a Sciex tripleTOF mass spectrometer using electrospray ionization. The compound was detected in positive and negative ion mode. Three microliters of samples were flow-injected by the autosampler onto a Waters Acquity UPLC BEH Amide column (1.7 μm, 2.1 × 100 mm). The mobile phase components consisted of (A) 20Mm Ammonium acetate, 20Mm Ammonium hydroxide and (B) CH3CN. The gradient profile used is as detailed as following: initial time, 5% A and 95% B; 0.5 min, 5% A and 95% B; 8 min, 35% A and 65% B; 9.5 min, 60% A and 40% B; 10.5 min, 60% A and 40% B; 11.5 min, 5% A and 95% B; 15.1 min, 5% A and 95% B. The flow rate was 0.45 mL/min. The mass spectrometry settings were as follows: Ion Source Gas 1 (GS1), 60; Ion Source Gas 2 (GS2), 60; Curtain Gas (CUR), 30; Temperature, 500; IonSpray Voltage, 5.5 kV; Collision Energy, 40; Data collection and analysis was performed using Sciex software.

### Intracellular ROS measurement

The intracellular ROS were determined by carboxy-H2DCFDA (Aladdin, H131224-50mg, Shanghai). RBCs were washed with cold PBS (PH=7.0) and incubated with 10uM carboxy-H2DCFDA in the dark for 30 min at 4 c. Intracellular fluorescent products were measured immediately by flow cytometry.

### Erythrocyte oxidation parameters detection

Erythrocyte ghosts were prepared according to a modification technique (Hoffman 1962). In short, the hemolysis of RBCs occurs in the hypotonic solution, and ghosts can be obtained by centrifugation to remove hemoglobin and inclusions. Sulfhydryl content in erythrocyte was measured by ELLMAN method (Smith*, et al* 1988). The level of carbonyl group in protein was determined by DNPH (2,4-Dinitrophenylhydrazine) (Reznick and Packer 1994). MAD (Malondialdehyde) level is quantified with a MAD kit (Jianchen Bioengineering Institute, A003-1-2, Nanjing).

## Results

### *Ggct* deletion in mice leads to splenomegaly and progressive anemia

*Ggct* deficient mice were generated through embryonic stem cell targeting and blastocyst injection as described previously (He*, et al* 2019). Homozygous *Ggct* knock-out mice were genotyped by PCR (Figure S1A). The depletion of Ggct protein was confirmed by Western blot analysis (Figure S1B). Although the *Ggct^−/−^* mice are viable and appear healthy, they were found to have splenomegaly (Fig. 1A). Spleen was 1.8 times larger (Fig. 1B) in the *Ggct^−/−^* mice than in the wild-type sibling controls. No significant difference in body weight was observed between *Ggct^−/−^* and the control littermates (He*, et al* 2019). Furthermore the weights of other major organs in *Ggct^−/−^* mice were not significantly changed compared to sibling controls (Figure S2). Hematoxylin and eosin-stained sections of spleens from *Ggct^−/−^* mice revealed expanded red pulps (Fig. 1C and D). Giemsa staining of blood smears indicated dysmorphic red cells, such as stomatocytes in *Ggct^−/−^* mice (Fig. 1E and F).

**Figure 1.**
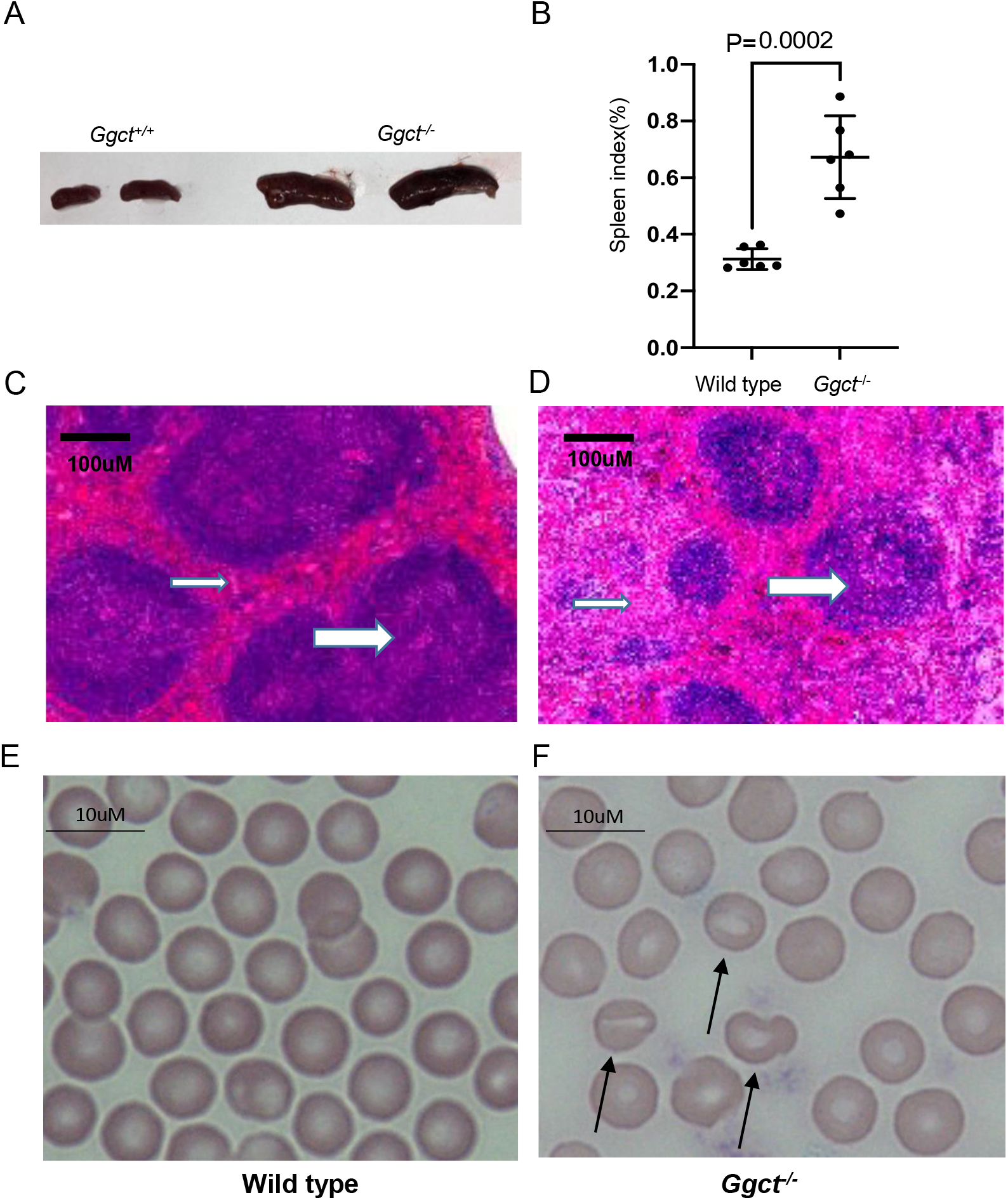
*Ggct* deletion in mouse leads to splenomegaly. (A)Representative images of six-month-old wild-type and *Ggct^−/−^*mouse spleen. (B) Spleen index (spleen weight/body weight) in wild-type and *Ggct^−/−^* mice. P value of Student’s t test is shown. (C-D) Hematoxylin and eosin-stained sections of spleen from wild-type (C) and *Ggct^−/−^* (D) mice. *Ggct^−/−^* mice show enlarged red pulp. Thick arrow indicates white pulp and thin arrow indicates red pulp. (E-F) Giemsa-stained blood smears from wild-type (E) and *Ggct^−/−^* (F) mice. Arrows indicate stomatocyte.

Hematological tests showed anemia in *Ggct^−/−^* mice, as evidenced by low erythrocyte number, low hemoglobin content, low hematocrit and increased reticulocyte count in *Ggct^−/−^* mice compared with wild-type mice (Fig. 2A-C, 2E). Meanwhile leukocyte counts were similar in wild-type and *Ggct^−/−^* mice (Figure S3). The anemia phenotype is more severe in adult or older *Ggct^−/−^* mice, and the RBC differences are not significant in very young (<3 months old) *Ggct^−/−^* mice compared to sibling controls (Figure S4). Collectively, these data indicate that *Ggct* deficiency in mouse leads to progressive anemia phenotype.

**Figure 2.**
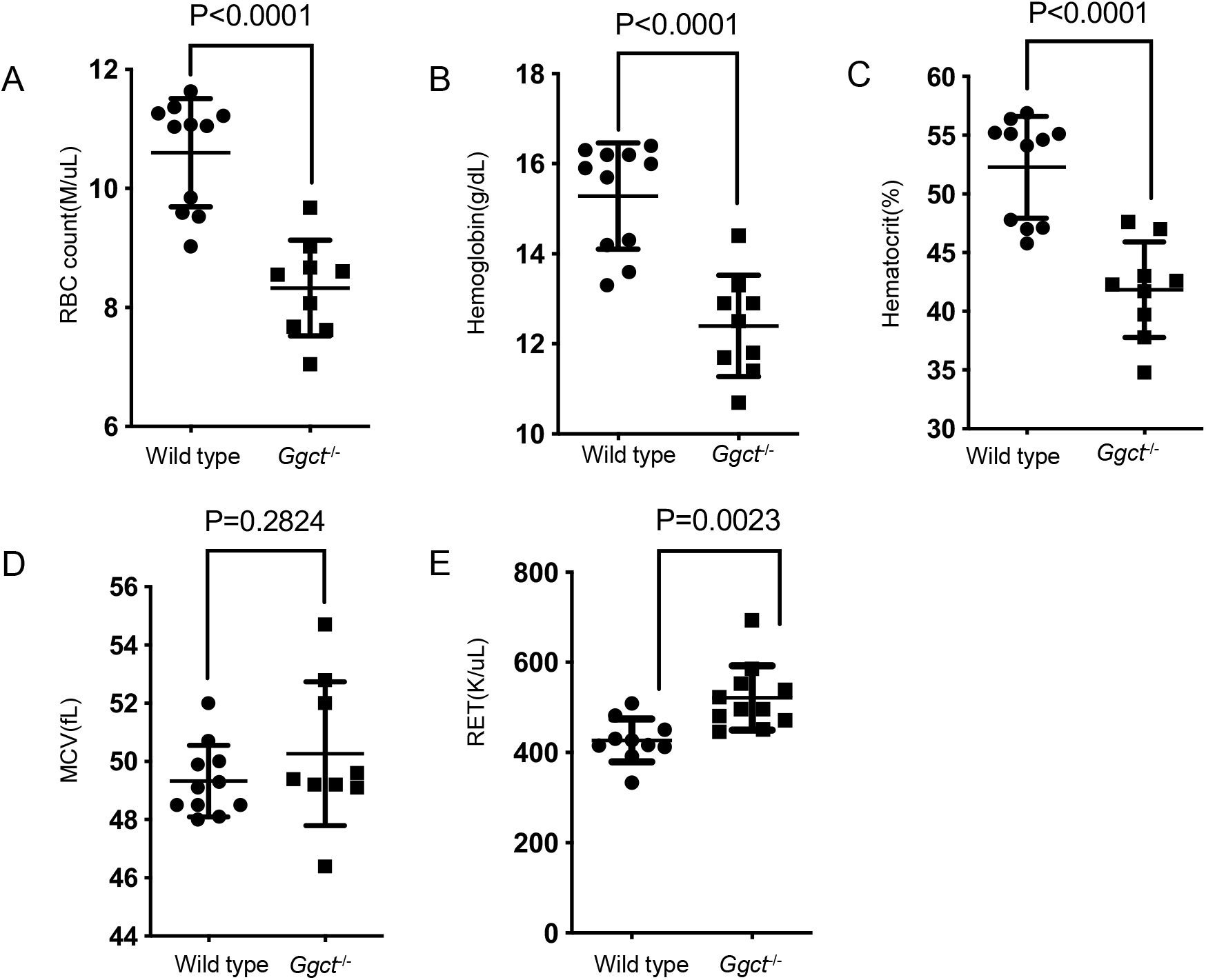
*Ggct^−/−^* mice show anemia phenotype. Hematologic parameters of six-month-old wild-type and *Ggct^−/−^* mice are shown. (A) RBC, red blood cells; (B) HGB, hemoglobin; (C) HCT hematocrit; (D) MCV mean cell volume; (E) RET reticulocytes. P values of Student’s t test are shown. Error bars indicate mean ± s.d.

### Elevated erythropoiesis in *Ggct^−/−^* mice

The anemia phenotype observed in *Ggct^−/−^* mice could be due to defective RBC maturation (ineffective erythropoiesis), increased RBC destruction (hemolysis), or a combination of both processes. We analyzed splenocytes by flow cytometry. A subpopulation of Ter119+ cells was distinguished based on their expression of the transferrin receptor (CD71), which decreases with erythroblast maturation (Dong*, et al* 2011). Using flow cytometry, we found that the proportions of erythroid precursor cells (Ter119+CD71+) in *Ggct^−/−^* mouse spleens were 2-fold higher than that in wild-type spleen (Fig. 3A and B). In hematologic analysis, reticulocytes in *Ggct^−/−^* mice were higher than in wild-type mice (Fig. 2E). The increased reticulocyte levels were in line with elevated plasma Epo protein in *Ggct^−/−^* mice (Figure S5). In addition, the number of erythroid precursor cells was higher in the bone marrow of *Ggct^−/−^* mice than wild-type mice (Fig. 3C and D), and the percentage of Ter119+CD71+ cells was increased 1.2-fold in bone narrow of *Ggct^−/−^* mice, from 23.76% in wild-type mice to 28.2% in *Ggct^−/−^* mice. These data suggest that erythropoiesis was elevated in *Ggct^−/−^* mice, possibly reflecting a compensatory reaction to the anemia phenotype of *Ggct^−/−^* mice.

**Figure 3.**
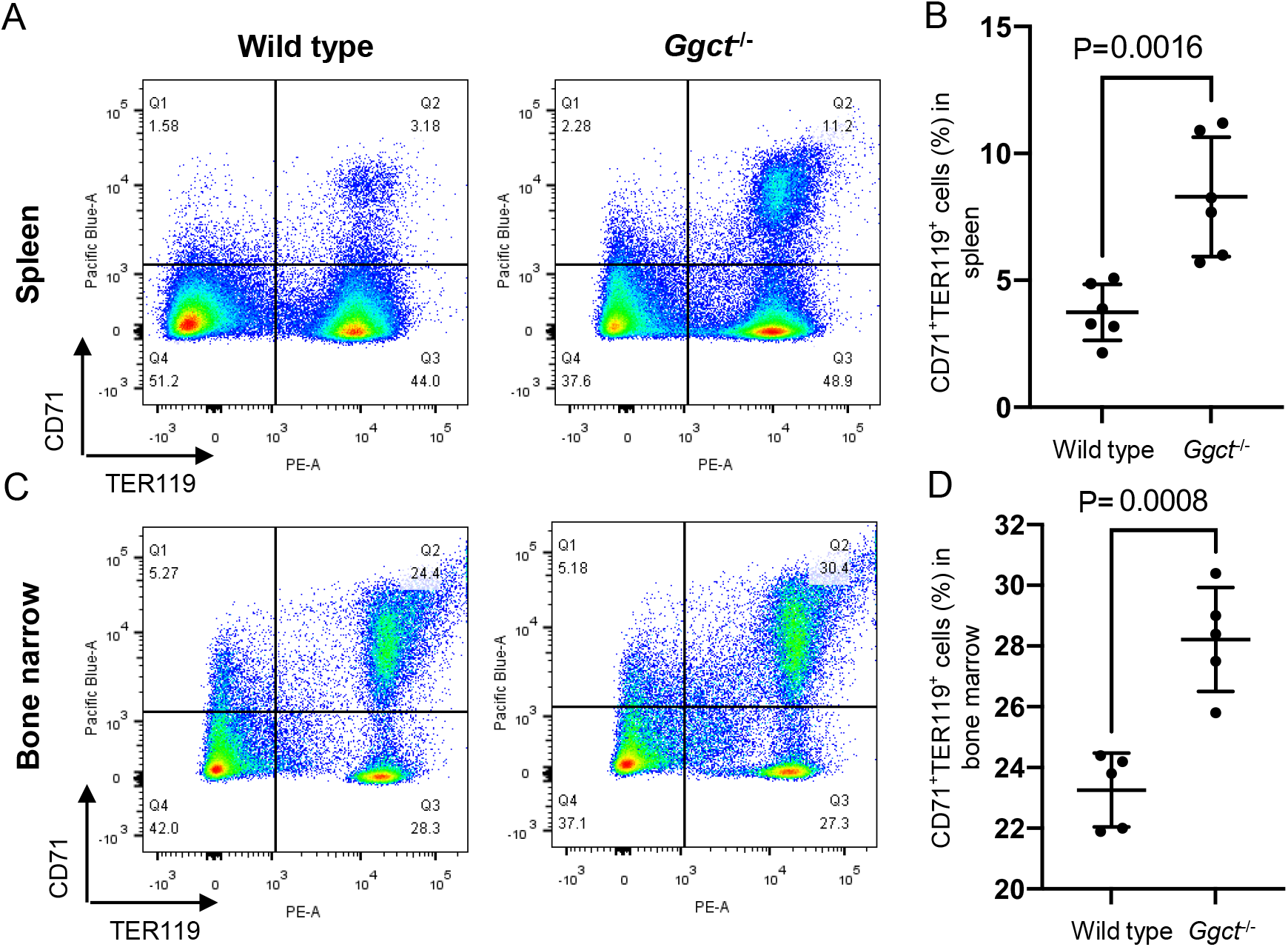
Elevated erythropoiesis in *Ggct^−/−^* mouse spleen and bone narrow. (A-B) Flow cytometry analysis of nucleated six-month-old mouse spleen cells, representative images (A) and statistical data (B) are shown. (C-D) Flow cytometry analysis of nucleated mouse bone marrow cells, representative images (C) and statistical data (D) are shown. P values of Student’s t test are shown. Error bars indicate mean ± s.d.

### Intracellular ROS levels and oxidation damages are increased in *Ggct^−/−^* RBCs

We measured intracellular reactive oxygen species (ROS) levels within RBCs with carboxy-H2DCFD fluorescent dye. Intracellular ROS concentrations measured under base-line conditions or after challenging with exogenous H_2_O_2_ (50uM) were elevated in *Ggct^−/−^* RBCs compared with wild-type RBCs (Fig. 4A and B). One of the typical features of damaged RBCs is the presence of lipid peroxidation and protein carbonylation. Protein oxidation can be measured by classic biochemical methods for carbonyls that result from the reaction of side chains of lysine, proline, threonine, or arginine with ROS. In agreement with the observed elevation in ROS concentrations, oxidized protein levels in *Ggct^−/−^* RBCs were markedly increased (1.5-fold) as measured by 2,4-dinitrophenylhydrazine-derivatized carbonyl (Fig. 4C). Owing to thiol groups are known to be easily oxidized by attack of ROS (Kim*, et al* 2000), we then monitored thiol group of cellular proteins by 5,5’-DiThiobis-2-NitroBenzoic acid (DTNB). The results show that the thiol group levels of membrane protein were decreased in *Ggct^−/−^* RBCs compared with wild-type RBCs (Fig. 4D). In addition, we found that lipid peroxidation measured by MAD (Malondialdehyde) content was up-regulated in *Ggct^−/−^* RBCs compared with wild-type RBCs (Fig. 4E). These data suggests that *Ggct^−/−^* RBCs are more susceptible to ROS, and suffer from serious oxidative damage, as evidenced by increased protein carbonyl groups and lipid peroxidation, decreased thiol groups in *Ggct^−/−^* RBCs compared with wild-type RBCs.

**Figure 4.**
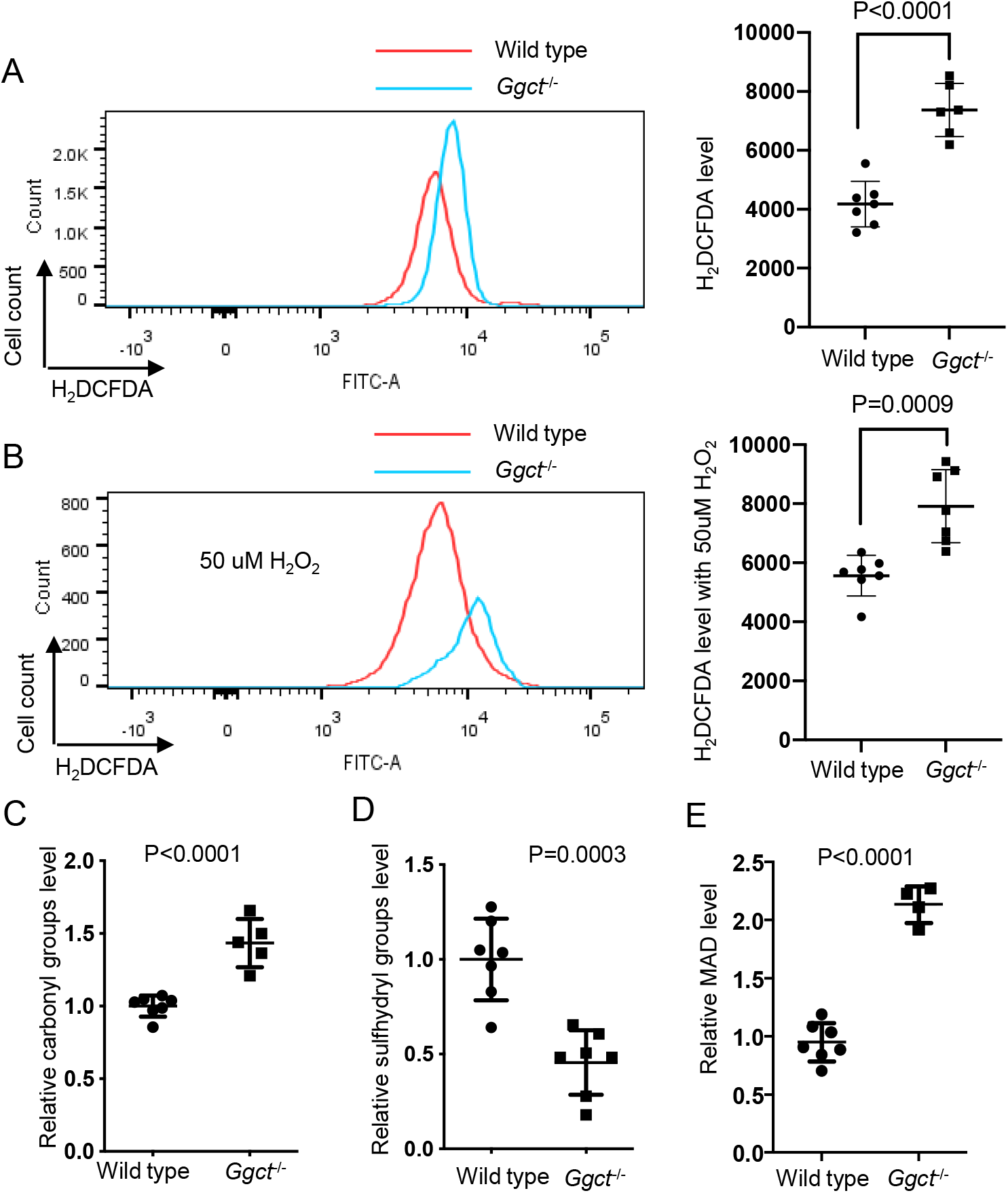
*Ggct^−/−^* RBCs show up-regulated ROS levels and oxidative damages. (A-B) Cellular ROS levels was determined in six-month-old wild-type and *Ggct^−/−^* RBCs by carboxy-H2DCFDA flow cytometry without (A) or with (B) the addition of exogenous H2O2 (50 uM). (C-E) The parameters of erythrocyte oxidative stress, including carbonyl groups (C), sulfhydryl group (D) and MAD level (E) in erythrocyte membrane proteins, were measured in wild-type and *Ggct^−/−^* mice. P values of Student’s t test are shown. Error bars indicate mean ± s.d.

### *Ggct^−/−^* RBCs show decreased life span

The normal life span of circulating RBCs is determined by their clearance from the peripheral circulation (predominantly by the spleen). We have shown that *Ggct* deficient erythrocytes are more susceptible to ROS; we therefore examined whether *Ggct* deficiency could affect RBC survival and whether the observed anemia was due to increased destruction of RBCs in the circulation *in vivo*. To this end, mice were infused with CMFDA labelled wild-type or *Ggct^−/−^* RBCs, and cell life span was determined by flow cytometric analysis of circulating labeled RBCs (Sandoval*, et al* 2008). We observed that labelled *Ggct^−/−^* RBCs disappeared more rapidly than wild-type RBCs (Fig. 5A), suggesting a faster clearance and a shorter lifespan of *Ggct^−/−^* RBC. To further demonstrate the destruction of erythrocytes in *Ggct^−/−^* mice, we performed a blood cross-transfusion experiment. Wild-type mice received either CMFDA-labeled wild-type erythrocytes (WT-WT) or *Ggct^−/−^* erythrocytes (KO-WT), and *Ggct^−/−^* mice received either CMFDA-labeled wild-type erythrocytes (WT-KO) or *Ggct^−/−^* erythrocytes (KO-KO). The result demonstrated that the KO-WT group had a higher clearance rate of infused erythrocytes than he WT-WT group, and similar results were observed when compared KO-KO with WT-KO group (Fig. 5B), suggesting an accelerated clearance of *Ggct^−/−^* erythrocytes. In line with the increased clearance rate, we observed that *Ggct^−/−^* RBCs were significantly less resistant to hypotonic lysis than wild-type RBCs (Fig. 5C).

**Figure 5.**
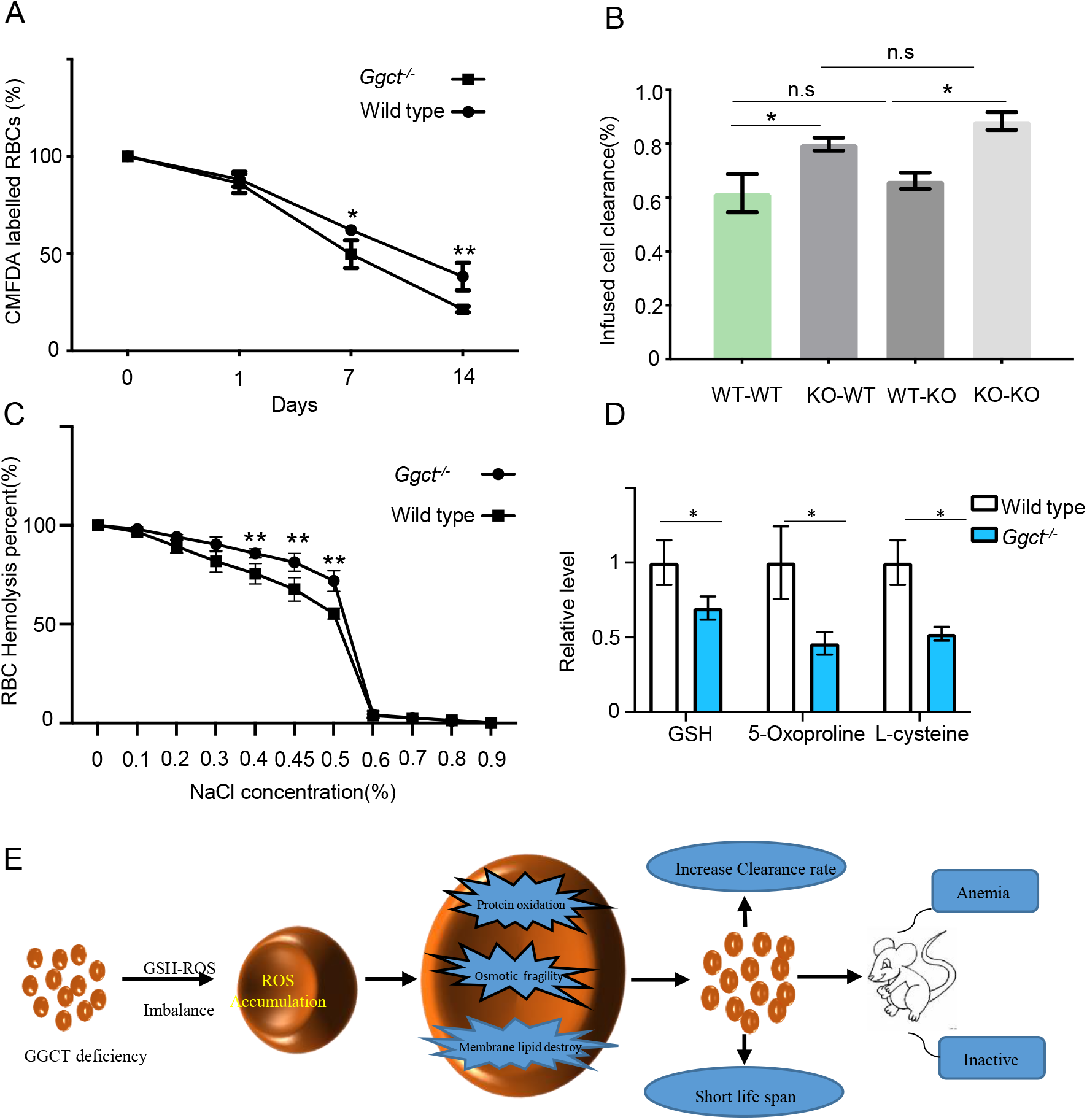
*Ggct^−/−^* RBCs show increased osmotic fragility and shortened life span. (A) Quantification of transferred CMFDA-labelled wild-type and *Ggct^−/−^* RBCs *in vivo*. (B) Clearance quantification of CMFDA-labeled RBCs 14 days after reinfusion. WT-WT, RBCs from wild-type donor were infused into wild-type recipients; KO-WT, RBCs from *Ggct^−/−^*donor were infused into wild-type recipients; WT-KO, RBCs from wild-type donor were infused *Ggct^−/−^* recipients; KO-KO, RBCs from *Ggct^−/−^* donor were infused into *Ggct^−/−^* recipients. (C) Osmotic fragility was quantified in wild-type and *Ggct^−/−^* mice (n=6 in each group). (D) Quantification of GSH metabolism related molecules in wild-type and *Ggct^−/−^* RBCs, error bars represent mean ± SD of three experiments. P values of Student’s t test are shown (*, *p*<0.05; **, *p*<0.01). (E) Proposed model for Ggct function in RBC. *Ggct* deficiency in RBC leads to the accumulation of ROS due to impaired GSH metabolism. Up-regulated ROS level decreased the lifespan of *Ggct^−/−^* RBCs. Splenomegaly and the lack of activity phenotype observed in *Ggct^−/−^* mice could be the consequence of anemia.

To explore the physiological consequence of *Ggct* loss induced anemia in mice, we performed metabolic cage experiments on *Ggct^−/−^* mice and wild-type mice. The results showed that the activity of *Ggct^−/−^* mice was significantly less than that of wild-type mice (Fig. S6A and B); particularly during the dark phase (Fig. S6C). Energy expenditure, measured as O2 consumption and CO2 production, was markedly reduced in *Ggct^−/−^* mice (Fig. S6D). Heat production, an indicator of metabolic rate, was also reduced significantly in *Ggct^−/−^* mice (Fig. S6F). Mice of both genotypes exhibited similar respiratory exchange ratio (RER) (RER = V_CO2_/V_O2_) (Fig. S6E), indicating that the loss of *Ggct* did not alter fuel preference.

### Metabolomics analysis in wild-type and *Ggct^−/−^* RBC

We compare the contents of small molecules in wild-type and *Ggct^−/−^* RBCs by metabonomics mass spectrometry. The results show that compared with wild-type mice, the content of GSH in RBCs is decreased, and the precursor molecule for GSH synthesis, such as L-cysteine is also reduced in *Ggct^−/−^* RBCs (Fig. 5D). 5-oxoproline is the reaction product of γ-glutamyl cyclotransferase enzyme activity, its concentration is also decreased in *Ggct^−/−^* RBCs (Fig. 5D). GGCT could affect cellular L-cysteine content through regulating amino acid transport during the glutathione cycle (Thompson and Meister 1976). L-cysteine is the rate-limiting substrate in glutathione synthesis (Lu 2013), its down-regulation can lead to the down-regulation of glutathione in *Ggct^−/−^* RBCs.

In together, our data indicate that *Ggct* deficiency affects the metabolic balance of GSH-ROS in RBCs, results in the up-regulation of ROS level, thus affects the life span and the physiological function of RBCs. Splenomegaly and the inactivity phenotypes observed in *Ggct^−/−^* mice could be due to the antioxidant defect of *Ggct^−/−^* RBCs (Fig. 5E).

## Discussion

In this study, we provide the first evidence to suggest that *Ggct* is required for RBC life span maintenance and antioxidant defense. The progressive anemia and inactivity phenotypes of *Ggct^−/−^* mice could be due to the defect in *Ggct^−/−^* RBC. This conclusion was supported by several lines of evidences. First, the rate at which labeled erythrocytes were eliminated from the circulation was markedly higher in *Ggct^−/−^* mice than in wild-type littermates (Fig. 5A). Consistent with this result, reticulocytes were increased in *Ggct^−/−^* mice (Fig. 2E), and the increased reticulocyte levels correlate with elevated plasma Epo protein (Fig. 3E). In addition, Ter119^+^CD71^+^ erythroblasts in the spleen and bone marrow were also markedly expanded (Fig. 2A-D).

Abnormal structure and deformability of the erythrocyte membrane plays an important role in shortened erythrocyte survival in many types of hemolytic anemia (An and Mohandas 2008). Consistent with this interpretation, we found the level of ROS in erythrocytes of *Ggct^−/−^* mice was up-regulated, resulting in the aggravation of erythrocytes oxidative damage. In accordance with this result, erythrocyte membrane proteins and lipids in *Ggct^−/−^* mice were oxidized, leading to a significant decrease osmotic fragility, an indication of increased rigidity of erythrocytes. As a result, the life span of erythrocytes in *Ggct^−/−^* mice is shorter than wild-type littermates. This phenotype is similar to that observed in mice deficient for the antioxidant enzymes, such as AMPKα1 (Wang*, et al* 2010), Nix (Sandoval*, et al* 2008) and prx II (Lee*, et al* 2003).

GSH is synthesized in the cytoplasm, and the availability of L-cysteine is the key determinants of GSH biosynthesis (Lu 2013). Before the identification of ChaC family proteins as the cytosolic pathway for glutathione degradation in mammalian cells (Kumar*, et al* 2012), GSH was thought to be degraded exclusively in the extracellular space by membrane-bound γ-glutamyl transpeptidase (GGT) to cysteinyl-glycine and γ-glutamyl–amino acid dipeptide (Ballatori*, et al* 2009). One of the best acceptor amino acids for GGT enzymatic reaction is L-cystine (Thompson and Meister 1976). In the absence of *Ggt*, intracellular GSH level is down-regulated due to decreased availability of intracellular L-cysteine (Bachhawat and Kaur 2017, Hanigan 2014). Based on our experimental data, *Ggct* deficiency also leads to decreased intracellular L-cysteine and consequently GSH down-regulation. Thus membrane bound Ggt and cytoplasmic Ggct could function together in GSH homeostasis through recycling L-cysteine.

In summary, using *Ggct^−/−^* mouse model, we demonstrate a critical function of *Ggct* in GSH metabolism, antioxidant defense and RBC life span maintenance. The progressive anemia and inactivity phenotypes of *Ggct^−/−^* mice could be due to the defects in *Ggct^−/−^* RBC.

## Author contributions

ZH, XS, DB, XZ maintained the mouse lines and performed the experiments; SW participated in critical project discussions; ZH, YH, PH performed the metabolomics experiments; XSL designed, supervised the study and wrote the manuscript.

