## Supplemental Figure for "Essential role for Ggct in erythrocyte antioxidant defense"

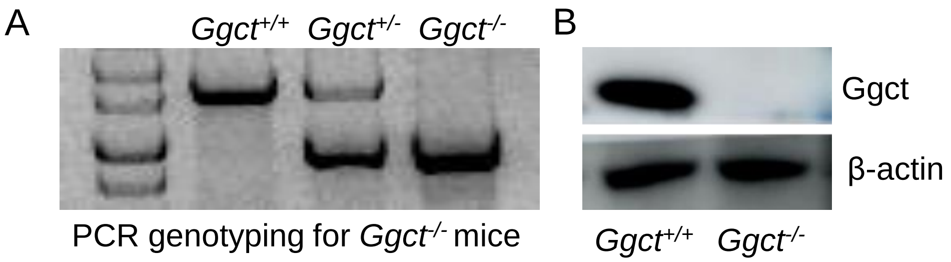


**Figure S1.** ***Ggct$^{-/-}$* mouse genotyping.** (A) PCR genotyping results of *Ggct* homozygous (-/-), *Ggct* heterozygous (+/-), and wild-type mouse (+/+). (B) Western blot analysis of *Ggct* expression in spleen from wild-type and *Ggct$^{-/-}$* mouse.

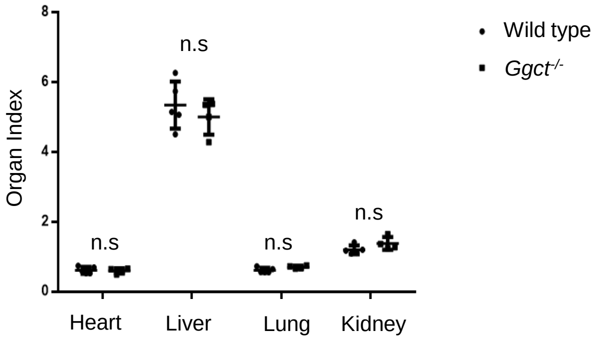


**Figure S2: Organ index.** No significant changes of heart index, liver index, lung index and heart index in *Ggct*$^{-/-}$ mice compared with wild-type mice. The data statistically analyzed by t test. Error bars indicate mean $\pm$ s.d.

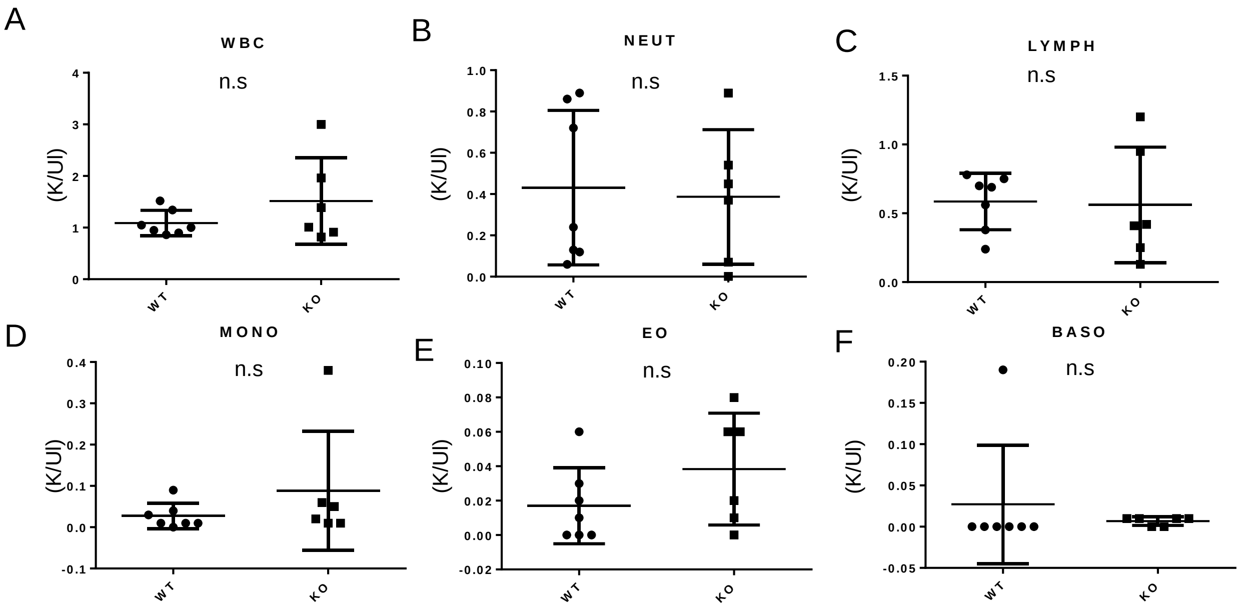


**Figure S3：Hematological parameters: analysis of leukocyte in mouse peripheral blood.** (A) WBC, white blood cell. (B) NEUT, neutrophil. (C) LYMPH, lymphocyte. (D) MONO, monocyte. (E) EO, eosinophilia. (F) BASO, basophil.

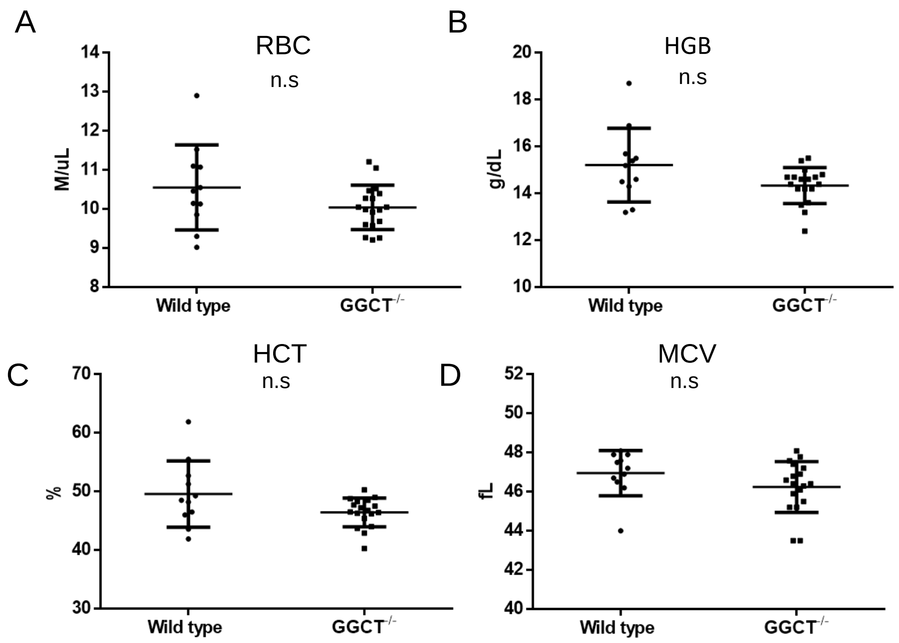


**Figure S4：Hematologic parameters of young (less than 15 weeks of age) *Ggct*$^{-/-}$ and wild-type mice.** (A-E) Selected blood routine parameters. (A) RBC, red blood cells; (B) HGB, hemoglobin; (C) HCT hematocrit; (D) MCV mean cell volume; The data statistically analyzed by t test.

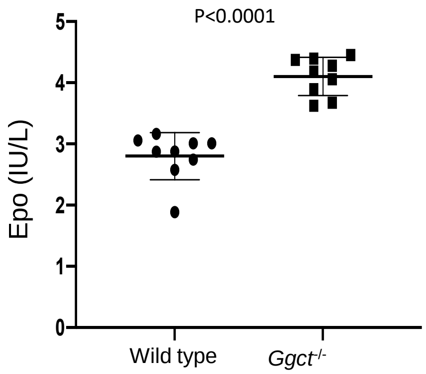


**Figure S5. Epo quantification in wild type and *Ggct$^{-/-}$* mouse.** Epo protein was quantified from wild-type and *Ggct$^{-/-}$* mouse plasma. P values of Student’s t test are shown. Error bars indicate mean $\pm$ s.d.

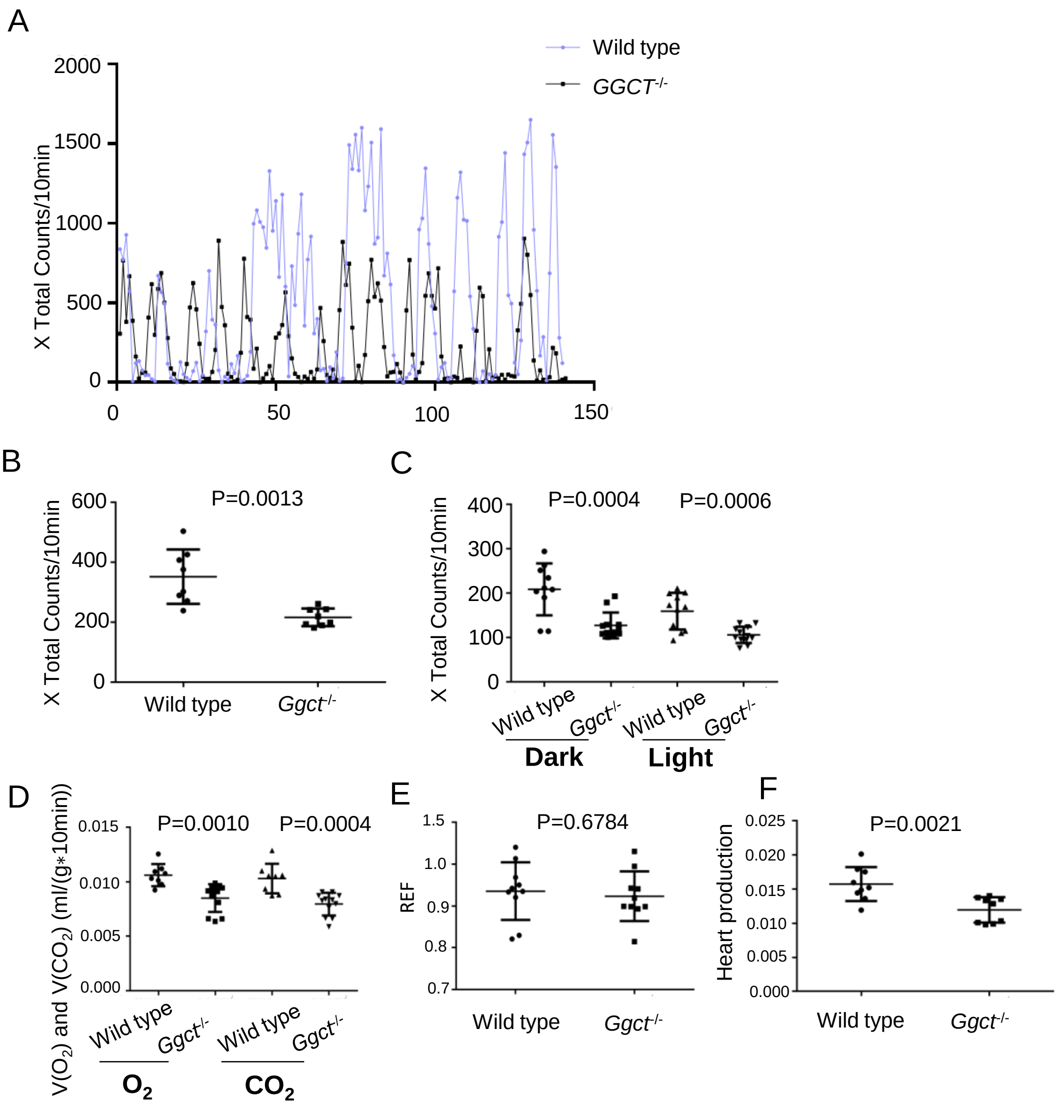
A
Wild type
GGCT-/-
X Total Counts/10min
2000
1500
1000
500
0
0
50
100
150
B
X Total Counts/10min
600
400
200
0
P=0.0013
Wild type
Ggct-/-
C
X Total Counts/10min
400
300
200
100
0
P=0.0004
P=0.0006
Wild type
Ggct-/-
Wild type
Ggct-/-
Dark
Light
D
V(O2) and V(CO2) (ml/(g*10min))
0.015
0.010
0.005
0.000
P=0.0010
P=0.0004
Wild type
Ggct-/-
Wild type
Ggct-/-
O2
CO2
E
REF
1.5
1.0
0.9
0.8
0.7
P=0.6784
Wild type
Ggct-/-
F
Heart production
0.025
0.020
0.015
0.010
0.005
0.000
P=0.0021
Wild type
Ggct-/-

**Figure S6: *Ggct*$^{-/-}$ mice show lack of activity phenotype.** (A) The activity of *Ggct* knockout mice was significantly lower than that of wild-type mice. Mouse movements were monitored in the metabolic chamber, and are quantified as the number of horizontal photobeam breaks per 10 min. (B) Statistical chart of mouse activity in (A). (C) The activity from wild type and *Ggct*$^{-/-}$ mice in dark and light was determined by the number of horizontal photobeam breaks per 10 min. (D) $O_2$ consumption ($V_{O2}$) and $CO_2$ production ($V_{CO2}$) were measured by Volume consumed per gram of mice per 10 minutes between wild type and *Ggct*$^{-/-}$ mice. (E) RER is calculated by $O_2$ consumption ($V_{O2}$) divide $CO_2$ production ($V_{CO2}$). (F) Heart production was determined by cage were recorded. All data were normalized to body weight, time and shown for a 24-hours period. P values of Student's t test are shown. Error bars indicate mean ± s.d. *P < 0.05; **P < 0.01; ***P < 0.001
